# Sex and Estrous Cycle in Memory for Sequences of Events in Rats

**DOI:** 10.1101/2021.10.29.466512

**Authors:** M. Jayachandran, P. Langius, F. Pazos Rego, R. P. Vertes, T. A. Allen

## Abstract

The ability to remember sequences of events is fundamental to episodic memory. While rodent studies have examined sex and estrous cycle in episodic-like spatial memory tasks, little is known about these biological variables in memory for sequences of events that depend on representations of temporal context. We investigated the role of sex and estrous cycle in rats during all training and testing stages of a cross-species validated sequence memory task (Jayachandran et al., 2019). Rats were trained on a task composed of two sequences, each with four unique odors delivered on opposite ends of a linear track. Training occurred in six successive stages starting with learning to poke in a nose port for ≥1.2s; eventually demonstrating sequence memory by holding their nose in the port for ≥1s for in-sequence odors and <1s for out-of-sequence odors in order to receive a water reward. Performance was analyzed across sex and estrous cycle (proestrus, estrus, metestrus, and diestrus), the latter being determined by the cellular composition of a daily vaginal lavage. We found no evidence of sex differences in asymptotic sequence memory performance, similar to published data in humans performing the analogous task (Reeders et al., 2021). Likewise, we found no differences in performance across the estrous cycle. One minor difference was that female rats tended to have slightly longer poke times, while males had slightly more short poke times but this did not affect their decisions. These results suggest sex and estrous cycle are not major factors in sequence memory capacities.

## INTRODUCTION

Episodic memory is the ability to encode and recollect our daily personal experiences (Tulving, 2002; Allen & Fortin, 2013). A fundamental feature of episodic memory is the ability to remember sequences of events as they occurred across time (Allen et al., 2020; Allen et al., 2014, 2016; Eichenbaum & Fortin, 2005; Jayachandran et al., 2019). While rodent studies have examined sex and estrous cycle as variables in spatial and episodic-like memory tasks, little is known about these biological variables in a sequence memory task that depends on representations of temporal contexts. The study of sex and sex hormones in sequence memory is particularly important in understanding the neurobiology of mental health disorders because of known sex differences and the prevalence of deficits in temporal cognition (Li & Singh, 201;4 Postma et al., 2004; Valera et al., 2010) in disorders such as schizophrenia and Alzheimer’s disease (Kessler et al., 2005; Launer, 1999; Pigott, 2003).

Sex differences in memory have been widely reported in human and nonhuman animals alike (Levy et al., 2005; Seymoure et al., 1996; Williams et al., 1990). For example, rodent studies have shown that sex impacts both spatial cognition and object recognition (Barha et al., 2010; Hamson et al., 2016; Koss & Frick, 2017; Luine et al., 2017; Spritzer et al., 2008). Most commonly, evidence suggests that human and rodent males have an advantage in tests of spatial memory (Driscoll, 2005; Jonasson, 2005; Nowak et al., 2014; Steimer & Woolley et al., 2010). Importantly, when strategy is considered or a landmark, such as a wall cue is included, these sex differences disappear (Chai & Jacobs, 2010; Saucier et al., 2008). Alternatively, females tend to outperform males in object memory (Bettis & Jacobs, 2012; Levy et al., 2005; Sutcliffe et al., 2007). However, not every study finds sex differences in spatial learning and memory tasks (Bucci et al., 2021; Frick et al., 1999; Healy et al., 1999).

When looking at sex differences in memory, it is important to consider the bioavailability of key sex hormones. For example, exogenous ovarian hormones have been shown to enhance performance in object memory in in both rodent and human females (Walf & Frye, 2006). This suggests the sex differences in memory can be dependent on the levels of circulating gonadal hormones. Important to our experiments here, hormone levels change reliably during the estrous cycle in females and different phases have been linked to differences in memory. For example, Healy and colleagues, (1999) showed that female rodents were slowest to locate the platform in a Morris water maze task during estrus and were fastest during proestrus which was attributed to a core spatial memory difference. Likewise, in other spatial tasks, females demonstrate the use of allocentric strategies during proestrus and egocentric/mixed strategies during diestrus (Korol, 2004; Koss & Frick, 2017). In object memory tasks, rats in proestrus outperformed rats in diestrus or metestrus (Koss & Frick, 2017; Van Goethem et al., 2012). By contrast to these aforementioned studies not all studies find differences in memory across the estrous cycle. For example, Stackman et al., (1997) did not observe significant differences in acquisition or performance levels across the estrous cycle when testing working memory using a radial arm maze; however, they showed that rats performed the task significantly slower during proestrus than on any other estrous cycle phase. This suggests that while the underlying working memory process remained stable, other aspects of performance can vary across the estrous cycle.

Here, we investigated the role of sex and estrous cycle (proestrus, estrus, metestrus, diestrus) in rats during training and testing stage of an odor sequence memory task (Jayachandran et al., 2019). Rats were trained for several months on the sequence memory task composed of two four-odor sequences delivered at opposite ends of a linear track. Training occurred progressively across six major stages starting with learning to poke and hold for ≥1.2s for a small water reward to eventually differentiating in-sequence (InSeq) and out-of-sequence (OutSeq) odors, one at a time, in four odor sequence sets. Performance across training stages was analyzed across sex and estrous cycle phases, the latter of which was determined by the cellular composition of a vaginal lavage. We conducted an extensive analysis of behavior at each stage including trials to criterion, self-paced inter-odor interval times, self-paced inter-sequence interval times, poke times under different sequential and accuracy conditions, and overall sequence memory. We found no evidence that males and females differed in learning, remembering, or performing the sequence memory task. Moreover, we did not find any differences across the estrous cycle in sequence memory. The results suggest sex and estrous cycle are not major factors in memory for sequences of events.

## METHODS

### Subjects

Twenty Long-Evans rats (10 females; Charles River Laboratories; weighing 250-350g upon arrival) were used. Two male rats failed to progress past the TwoOdor stage of training and were excluded from all analysis. Rats were individually housed and maintained on a 12-hr inverse light/dark cycle (lights were turned off at 10 AM). Rats had *ad libitum* access to food, but access to water was limited to 3 - 5min each day, depending on how much water they received as a reward during behavioral training (6 - 9mL). All training sessions and lavages were conducted during the dark phase (active period) of the light cycle. All experimental procedures using animals were conducted in accordance with the Florida International University Institutional Animal Care and Use Committee (FIU IACUC).

### Task Apparatus

Rats were tested in a noise-attenuated experimental room. The behavioral apparatus consisted of a linear track (length, 183cm; width, 10cm; height, 43cm) with walls angled outward at 15° and nose ports centered at each end through which repeated deliveries of multiple distinct odors could be presented. Photobeam sensors were used to detect nose port entries. Each nose port was connected to an odor delivery system (Med Associates; Fairfax, VT). Odor deliveries were initiated by a nose-poke entry and terminated either when the rat withdrew their poke or after 1s had elapsed. Water ports were positioned under each nose port for reward delivery. Timing boards (Plexon; Dallas, TX) and digital input/output devices (National Instruments; Austin, TX) were used to measure all event times and control the hardware. All aspects of the task were automated using custom MATLAB scripts (MathWorks 2021a; Natick, MA). A 256-channel OmniPlex with video tracking and CinePlex behavioral software (Plexon; Dallas, TX) were used to interface with the hardware in real time and record behavioral data. Odors were organic odorants contained in glass jars (Sigma-Aldrich; St. Louis, MO, A1: 1-octanol, CAS: 111-87-5; B1: (-) - limonene, CAS: 5989-54-8; C1: I-menthone, CAS: 14073-97-3; D1: isobutyl alcohol, CAS: 78-83-1; A2: acetophenone, CAS: 98-86-2; B2: (1S) - (-) – beta-pinene, CAS: 18172-67-3; C2: L (-) - carvone, CAS: 6485-40-1; D2: 5-methyl-2-hexanone, CAS: 110-12-3) that were volatilized with nitrogen air (flow rate, 2L/min) and diluted with ultrapure air (flow rate, 1L/min). To prevent cross-contamination, separate Teflon tubing lines were used for each odor. These lines converged into a single channel at the bottom of the odor port. In addition, a vacuum located at the top of the odor port provided constant negative pressure to quickly evacuate odor traces with a matched flow rate.

### Sequence Memory Task

The sequence memory task (Jayachandran et al., 2019; Allen et al., 2014) involves repeated presentations of odor sequences and requires the rat to determine whether each item (odor) presented is in-sequence (InSeq) or out-of-sequence (OutSeq). Rats were trained on two sequences, each comprised of four distinct odors (e.g., Seq1: A_1_B_1_C_1_D_1_, Seq2: A_2_B_2_C_2_D_2_). Each sequence was presented at either end of a linear track maze. Odor presentations were initiated by a nose-poke, and each trial was terminated after the rat either held the nose-poke response for ≥1s (InSeq) or withdrew its nose-poke response <1s (OutSeq). Trials were separated by 1s intervals. Water rewards (20μl; diluted at 1g of aspartame for every 500mL of water) were delivered below the odor port after each correct response. Incorrect responses triggered a buzzer sound and the termination of the sequence. Each sequence was presented 50-100 times per session; approximately half the presentations included all items InSeq (ABCD) and half included one item OutSeq (e.g., AB<u>A</u>D, odor A repeated in the 3rd position). Note that OutSeq items could be presented in any sequence position except the first position (i.e., sequences always began with an InSeq item). Sequence memory was probed with OutSeq trials (e.g., AB<u>A</u>D; one OutSeq trial randomly presented per sequence).

### Sequence Memory Task Training

Naive rats were initially trained on a series of incremental stages over 20-30 weeks. Each rat was trained to poke and hold its nose in an odor port to receive a small water reward. The minimum required nose-poke duration started at 50ms and was gradually increased (in 15ms increments) until the rat held the nose-poke position for ≥1.2s ~ 75% of the time over three sessions (75-100 nose-pokes per session). The rats were then trained to poke on side 2 until reaching criterion; poking for ≥1.2s with ~75% accuracy for three consecutive sessions. The rats were then habituated to odor presentations in the port (odor A_1_ and A_2_). The hold time was reduced down to 1 s and rats were required to alternate between sides for a small water reward at each side for ~75% accuracy for three consecutive sessions before moving on. Then a second set of odors was introduced (Seq1: A_1_B_1_ and Seq2: A_2_B_2_) and each rat was required to maintain its nose-poke response for ≥1s to receive a reward and needed to achieve ~70% accuracy before moving on. The rats were next trained to identify InSeq and OutSeq items. Rats began this stage trained on a two-item sequence: they were presented with ‘‘AB’’ and “A<u>A</u>” sequences. The correct response to the first odor was to hold their nose-poke for ≥1s (Odor A always being the first item). For the second odor, rats were required to determine whether the item was InSeq (AB; hold for ≥1s to receive reward) or OutSeq (A<u>A</u>; withdraw before 1s to receive a reward). After reaching criterion on the two-item sequence, the number of items per sequence was increased to three odors each side (Seq1: A_1_B_1_C_1_ and Seq2: A_2_B_2_C_2_), and four odors each side (Seq1: A_1_B_1_C_1_ D_1_ and Seq2: A_2_B_2_C_2_D_2_) in successive stages. The criterion for these stages set percentages as indicated in the results or the ability differentiate InSeq and OutSeq items above a sequence memory index (SMI; see quantitative and statistical analysis) of 0.2 over three consecutive sessions. Once rats reached criterion for the last stage of training, 15 consecutive sessions were collected to examine both sex and estrous cycle differences in overall SMI.

### Lavaging

Phases of the estrus cycle were determined in each female rat using vaginal lavage within four hours of each behavioral session. Rats were restrained using a designated towel and a drop of sterilized saline (NaCl, 0.85%) was placed over the vaginal opening to clean the area. The tip of a sterilized transfer pipette filled with sterilized saline (~0.25 – 0.5mL) was inserted into the vaginal opening to extract vaginal cells. All vaginal smears were placed on glass slides and photomicrographs were taken under brightfield illumination using an Olympus BX41 following guidelines from Westwood (2008). To control for experimenter handling, male rats were handled similarly and poked around the anal opening with a clean transfer pipette.

### Estrous Cycle Cytology

Estrous cycles in rats typically last four to five days and have four distinct phases: proestrus, estrus, metestrus and diestrus; each phase corresponds with specific cell types, i.e., nucleated epithelial cells, amorphous cornified cells, and small round leukocytes. Proestrus is characterized predominantly by nucleated epithelial cells, estrus predominantly amorphous cornified cells, metestrus by a mixture of nucleated epithelial cells, cornified cells and leukocytes, and diestrus predominantly leukocytes. All vaginal cell samples were obtained and imaged, then the images were blindly classified by three investigators.

### Quantification and Statistical Analysis

All data was analyzed in MATLAB 2021a (MathWorks; Natick, MA), SPSS 20.0.0, and Excel 2016 using custom scripts and functions. Performance on the task can be analyzed using a number of measures (Allen et al., 2014). The first position of each sequence was excluded from all analysis as these items are always InSeq. Expected vs. observed frequencies were analyzed with G tests to determine whether the observed frequencies of InSeq and OutSeq responses for a given session were significantly different from the frequency expected by chance. G tests provide a measure of performance that controls for response bias and is a robust alternative to the χ^2^ test, especially for datasets that include cells with smaller frequencies (Sokal & Rohlf, 1995). To compare performance across sessions or animals, a SMI was calculated (Allen et al., 2014) as shown in the following equation:

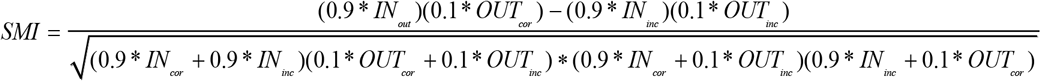

The SMI normalizes the proportion of InSeq and OutSeq items presented during a session and reduces sequence memory performance to a single value ranging from −1 to 1. A score of 1 represents a perfect sequence memory performance in which a subject would have correctly held their nose-poke response to all InSeq items and correctly withdrawn on all OutSeq items. A score of 0 indicates chance performance. Negative SMI scores represent performance levels below that expected by chance. SMI was calculated for males and females as well as between estrous phases during the two, three, and four odor stages of the sequence task. Using SPSS, an independent t-test was used to determine if males and females differed in their overall SMI. We went on to examine the differences between sequence 1 and sequence 2 within sex and estrous cycle by running a two-way ANOVA. A one-way ANOVA was used to determine if overall SMI differed within female estrous cycle. Linear regressions were performed in increments of pre-determined set trials or sessions to compare males and females in order to determine learning at each stage of training on the sequence task. Independent sample t-tests were used to analyze whether males and females differed in self-paced inter-trial-interval (ITI), inter-odor-interval (IOI), and inter-sequence interval (ISI). Nose-poke duration was analyzed using independent t-tests to determine whether rats held their responses significantly longer in InSeq_correct_ than in OutSeq_correct_ trials. General poke distributions were created through MATLAB using the session data.

## RESULTS

### Overall Sequence Memory Task and Training

We trained male (*n* = 10) and female (*n* = 10) rats in an odor sequence memory task (Jayachandran et al., 2019). The behavioral setup was automated and allowed repeated delivery of pure chemical odorants between two odor ports located at opposite ends of a linear track (Figure1A). The rats were trained to indicate if an odor is InSeq by holding their nose in the nose-port for ≥1s (Figure 1Bi) or OutSeq by withdrawing their nose prior to 1 s for a small water reward (Figure 1Bii). Rats were trained in progressive stages over several months on the sequence task, as depicted in Figure 1C. At each stage of training on the sequence task, rats had to reach a specific performance criterion in order to progress to the next stage. Once they reached the final stage (FourOdors) and behavioral criterion, we collected 15 consecutive sessions to analyze overall sequence memory. Two male rats were not able to continue past TwoOdors and therefore we removed them from the overall analysis. We calculated an SMI to measure overall sequence memory while controlling for individual differences in poke-hold behavior (Allen et al., 2014). We averaged the overall number of sessions per stage of training between males (Poke&Hold_Side1_: 28.125 ± 2.601; Poke&Hold_Side2_: 8.625 ± 1.731; OneOdor: 15.500 ± 2.706; TwoOdors: 49.375 ± 8.268; ThreeOdors: 33.500 ± 8.203; FourOdors: 30.625 ± 1.499) and females (Poke&Hold_Side1_: 22.800 ± 1.504; Poke&Hold_Side2_: 8.400 ± 1.962; OneOdor: 13.300 ± 1.506; TwoOdors: 40.600 ± 4.861; ThreeOdors: 30.800 ± 3.782; FourOdors: 30.400 ± 0.957). There were no significant differences in the number of sessions to criterion for stage when comparing males and females (Poke&Hold_Side1_: t_(16)_ = −1.861, *p* = 0.081; Poke&Hold_Side2_: t_(16)_ = −0.084, *p* = 0.934; OneOdor: t_(16)_ = −0.749, *p* = 0.465; TwoOdors: t_(16)_ = −0.959, *p* = 0.352; ThreeOdors: t_(16)_ = −0.320, *p* = 0.753; FourOdors; t_(16)_ = −0.131, *p* = 0.897).

**Figure 1.**
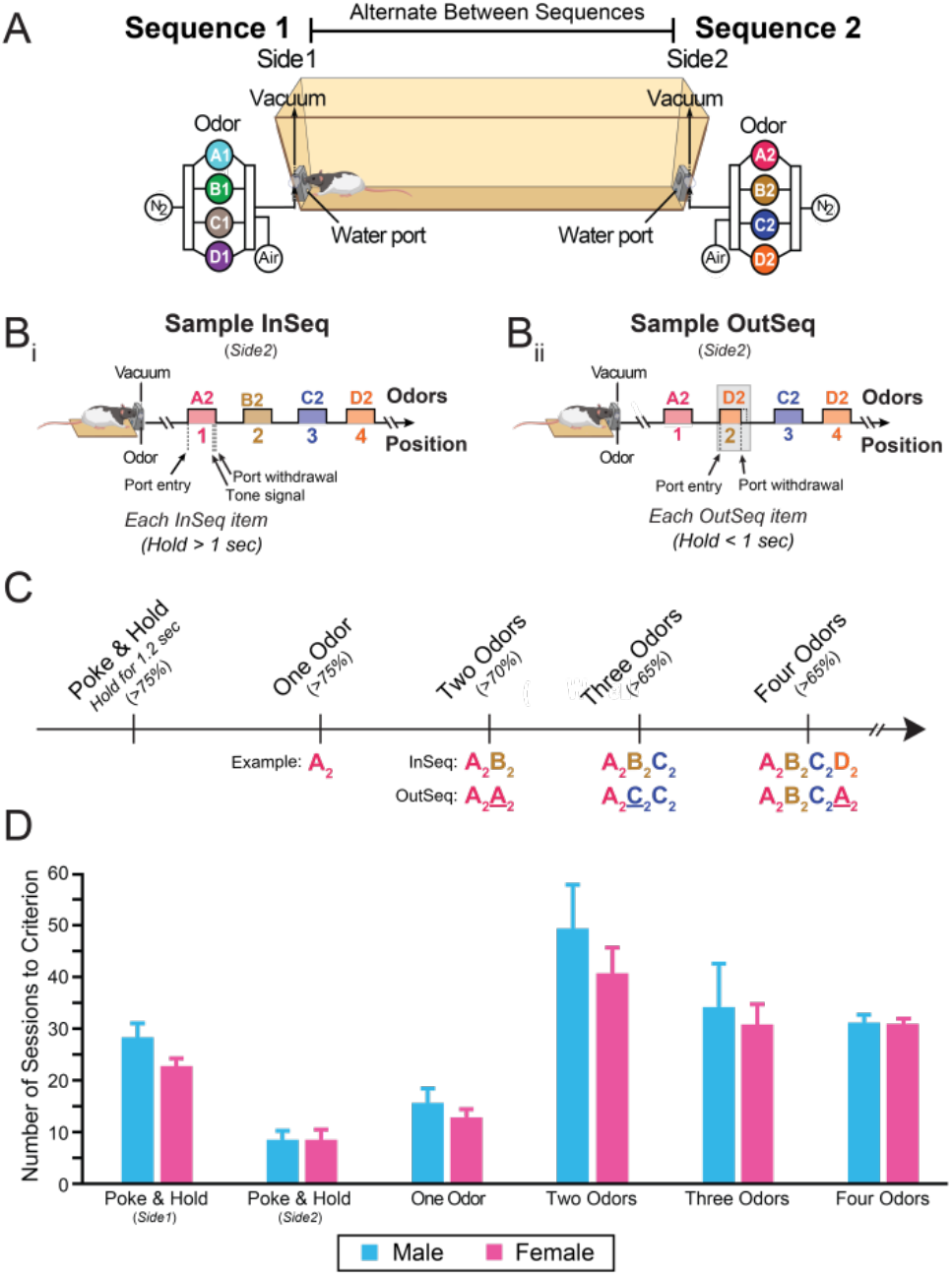
Sequence Memory Task. (A) A 2m linear track was used with odor ports located at opposite ends where two four-odor sequences were presented (Seq1: A_1_B_1_C_1_D_1_ and Seq2: A_2_B_2_C_2_D_2_). (B) Rats had to correctly identify the odors as either InSeq by holding their nose in the port for > 1 s (Bi) or OutSeq where the rat withdrew their nose prior to 1 s (Bii). (C) Timeline showing the criterion for each training stage for the sequence memory task. (D) Males and females did not differ in the average number of sessions per training stage. All data are represented as mean ± SEM. *p < 0.05; **p < 0.01; ***p < 0.001

### Males and Females Parallel in Poking Behaviors

Rats were first trained to poke and hold in the nose port located on side 1 of the sequence memory task (Figure 2Ai). During this stage, the rat is trained to hold their nose in the nose port with a start hold time of 50ms. With each correct trial, the hold time was increased by 15ms until reaching a 1.2s hold time. The rats were then required to poke for ≥1.2s at ~75% accuracy for 3 consecutive sessions. We first examined the learning rates of males and females by calculating the slope over 200 trial increments (Figure 2Aii). Males and females did not show any significant differences in learning during Poke&Hold_side1_ (Trials_1-200_: t_(16)_ = 1.075, *p* = 0.299; Trials_200-400_: t_(16)_ = 0.850, *p* = 0.262; Trials_400-600_: t_(16)_ = 0.336, *p* = 0.741; Trials_600-800_: t_(16)_ = 0.840, *p* = 0.413; Trials_800-1000_: t_(16)_ = 0.754, *p* = 0.462). A few rats exceeded 1000 trials to progress to the next stage of the task; however, due to too few values we could not calculate an average past 1000 trials. We then looked at the poke distributions between males and females for all trials. Overall, the proportion of nosepokes remained similar between males and females with a slight increase for males on short pokes (Figure 2Aiii). We performed several analyses to test the alternative hypothesis that non-memory-related behavioral effects account for impaired performance in the sequence task. We measured the time it took rats to initiate each poke (inter-trial-interval; ITI). The ITI did not differ significantly between males and females (Figure 2Aiv; t_(410)_ = −0.174, *p* = 0.862).

**Figure 2.**
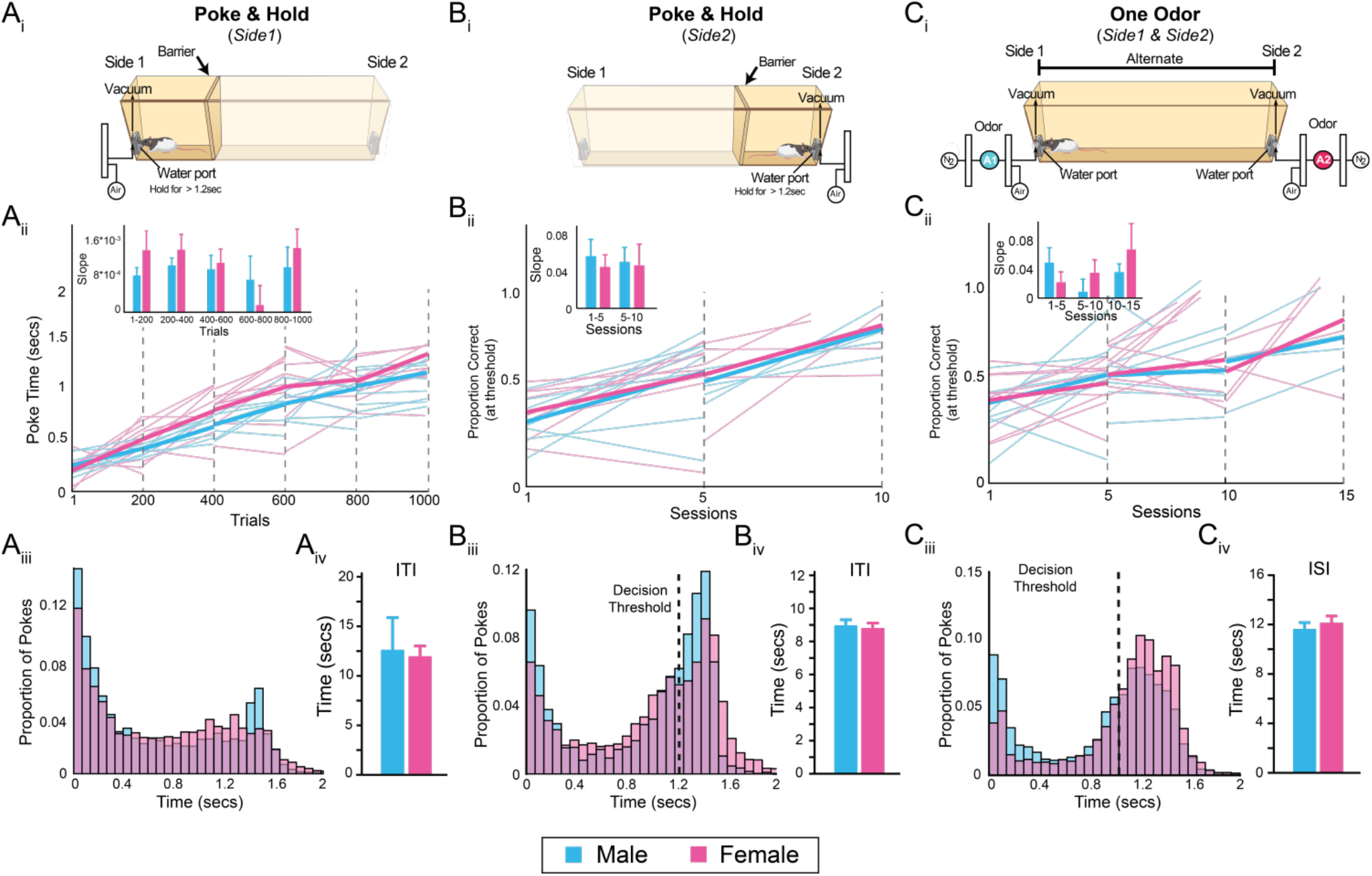
Males and Females Do Not Differ in Poking Behaviors. (A_i_) Rats were trained to hold their nose in the nose-port on side 1 for 1.2 s. (A_ii_) Males and females show no significant differences in the rate of learning across trials. (A_iii_) Males and females poke times were relatively similar. (A_iv_) Inter-trial-interval (ITI) was not significantly different between males and females (B_i_) Rats were trained to hold their nose in the nose-port on side 2 for 1.2 s. (B_ii_) No significant differences found between males and females in the rate of learning across sessions. (B_iii_) Males and females show subtle shifts in behavior where males show a tendency to have short poke times. Both males and females had increased number of pokes at the decision threshold (1.2 s). (B_iv_) ITI between sexes were not significantly different. (C_i_) Rats were introduced to one odor on either side of the maze and were trained to alternate holding their nose in the pose-port on both sides of the maze for 1 s. (C_ii_) No significant differences found between males and females in the rate of learning across sessions. (C_iii_) Male and female rats show subtle shifts in behavior where males show a tendency to have shorter poke times. (C_iv_) ISI between males and females was not significantly different. All data are represented as mean ± SEM. *p < 0.05; **p < 0.01; ***p < 0.001

Once reaching criterion for Poke&Hold_side1_ rats were then trained on Poke&Hold_side2_ where rats were familiarized with the odor port on side 2 and trained to poke for ≥1.2s at ~75% accuracy for 3 consecutive sessions (Figure 2Bi). We calculated the proportion correct for each session after poke hold times became consistent. We examined learning between males and females by measuring proportion correct and calculating the rate of change in proportion correct (Figure 2Bii; 5 session increments). A few rats continued past 10 sessions; however we were not able to calculate an average due to the low sample sizes after this point. Males and females did not show any significant differences in learning (Sessions_1-5_: t_(16)_ = −0.545, *p* = 0.593; Sessions_5-10_: t_(7)_ = −0.124, *p* = 0.905). When we looked at poke distributions, the proportion of nose-pokes between males and females remained similar (Figure 2Biii). However, we do note that males seem to have more short pokes (~ 0.1s) while females tended to hold longer (~1.4 s). We then examined ITI and found no significant differences between males and females (Figure 2Biv; t_(142)_ = −0.258, *p* = 0.797).

Once past Poke&Hold_side2_, rats were then moved to OneOdor which began with habituating to one odor presentation on either end of the linear track. The hold time criterion was also reduced down to 1 s and rats were required to alternate between sides (Figure 2Ci). Rats were required to reach ~75% accuracy for 3 consecutive sessions at this stage. We found no significant differences in learning between males and females (Sessions_1-5_: t_(16)_ = −1.123, *p* = 0.278; Sessions_5-10_: t_(16)_ = 1.034, *p* = 0.317; Sessions_10-15_: t_(7)_ = 0.774, *p* = 0.464). We then looked at poke distributions between males and females; overall, we found they were very similar. Again, we observed males’ tendencies towards short pokes (~0.1 - 0.2s) with females’ tendencies towards longer pokes (~1.2 - 1.5s). At this stage we measured the time it took the rats to run between sequences which we refer to as inter-sequence-interval (ISI). We found no significant differences in ISI between males and females (Figure 2Civ; t_(224)_ = 0.655, *p* = 0.513). Overall, these results suggest that males and females show little to no differences in their poking and timing behaviors.

### Males and Females Perform Similarly on the Sequence Memory Task

After reaching criterion for the OneOdor stage of training, rats are then moved onto TwoOdors where they are trained to poke for two separate odors on each side (Figure 3Ai; Seq1: A_1_B_1_ and Seq2: A_2_B_2_). Once rats are able to poke twice on each side with ~70% accuracy, they are then trained to identify InSeq and OutSeq odors. The correct response to the first odor is always to hold the nose-poke for ≥1s (Odor A was always the first item). For the second odor, rats were required to determine whether the item was InSeq (AB; hold for 1 s to receive reward) or OutSeq (A<u>A</u>; withdraw before 1s to receive a reward). Rats were trained until they demonstrated sufficient performance (≥ 0.2 SMI) for three consecutive sessions. We calculated the SMI once OutSeq trials were introduced and examined the overall learning between males and females. Although we refer to memory in this stage as sequence memory, it is more equivalent to a delayed-to-non-match sample task which exclusively measures working memory. Position 1 was excluded because an OutSeq item was never presented in that position. Twenty sessions were used to measure learning at this stage. While some rats continued to learn past 20 sessions, there were not enough subjects to calculate an average. Generally, we found no significant differences in learning between males and females (Sessions_1-5_: t_(16)_ = 1.140, *p* = 0.227; Sessions_5-10_: t_(16)_ = 0.685, *p* = 0.503; Sessions_10-15_: t_(15)_ = - 0.938, *p* = 0.363; Sessions_15-20_: t_(10)_ =-0.394, *p* = 0.702). We also examined the overall SMI of the 3 consecutive stages in which criterion was met. We found no significant difference between the sexes (Figure 3Aiii; t_(52)_ = 1.134, *p* = 0.262, Male: 0.347 ± 0.022, Female: 0.386 ± 0.026). Further, we examined whether there were differences between sequence 1 and sequence 2. A two-way ANOVA revealed no statistically significant interaction between sex and sequence (F_(3,104)_ = 0.031, *p* = 0.860). Simple main effect analysis showed that sex and sequence did not have a statistically significant effect on SMI (Sex: *p* = 0.304; Sequence: *p* = 0.643).

**Figure 3.**
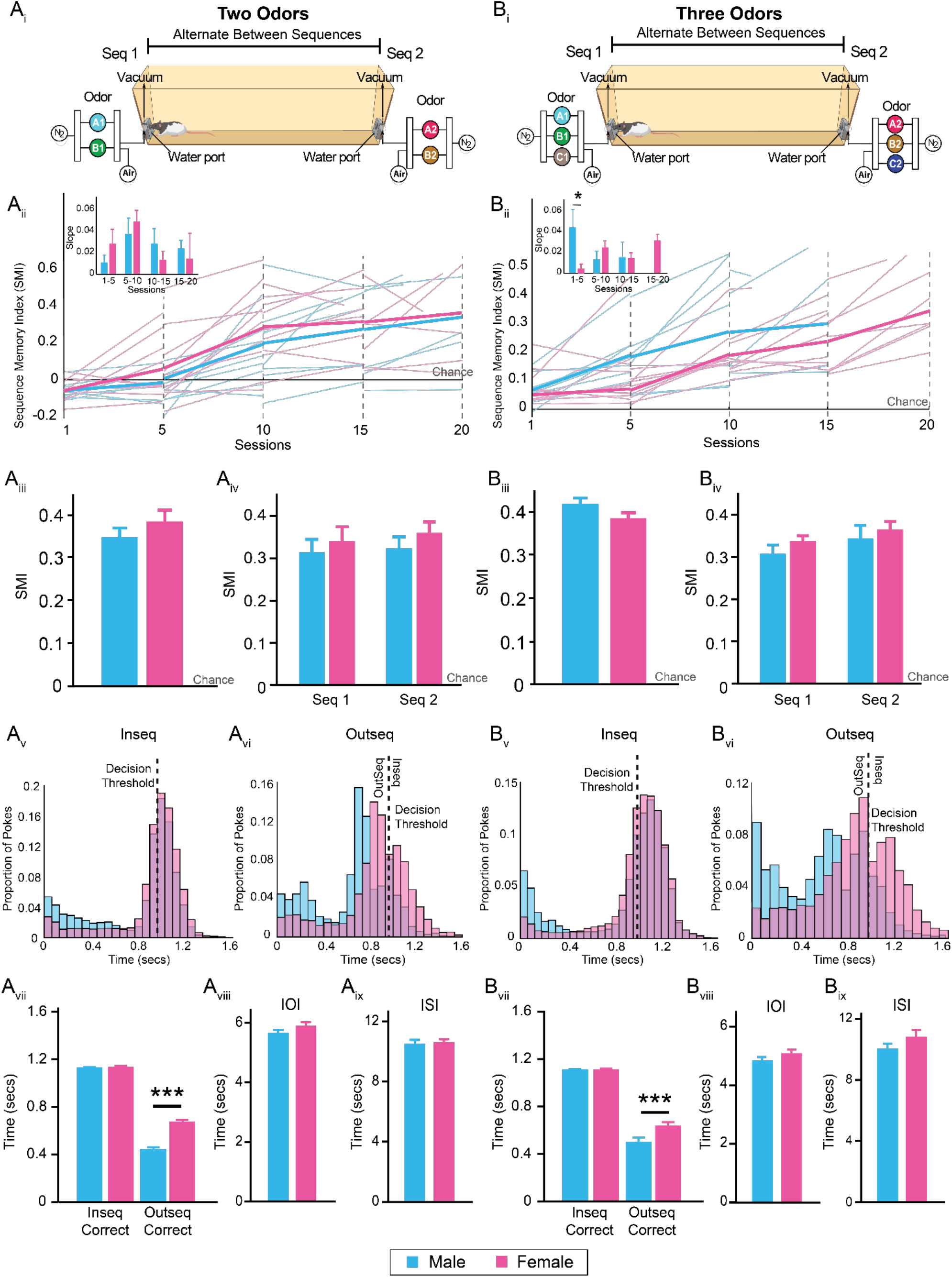
Learning Differs Between Males and Females. (A_i_) A second odor was added to either side of the maze (Seq1: A_1_B_1_ and Seq2: A_2_B_2_). (A_ii_) Performance did not differ between males and females in the rate of learning across sessions. (A_iii_) SMI was not significantly different between males and females. (A_iv_) SMI was not significantly different between sequence 1 and sequence 2 (A_v_) Males and females poke times were relatively similar. (A_vi_) Female nose-poke times show an increase in longer hold times while male poke times distribute between short and long poke times (A_vii_) InSeq_correct_ trials did not differ between males and females. OutSeq_correct_ nose-poke times were significantly different between males and females. (Av_iii_) Inter-odor-interval (IOI) was not significantly different between sexes (A_ix_) Inter-sequence-interval (ISI) was not significantly different between males and females (B_i_) A third odor was added to both sides of the maze (Seq1: A_1_B_1_C_1_ and Seq2: A_2_B_2_C_2_). (B_ii_) Learning rate between males and females was statistically significant during the first five sessions. As sessions continued, males and females no longer differed. (B_iii_) SMI was not significantly different between males and females. (B_iv_) SMI was not significantly different between sequence 1 and sequence 2. (B_v_) Male and female poke times for InSeq trials were relatively similar. Males have an increase in shorter poke times. (B_vi_) Males show an increase number of short pokes for OutSeq trials while females showed longer hold times. (B_vii_) InSeq_correct_ trials did not differ between males and females. OutSeq_correct_ poke times were significantly different between males and females. (B_viii_) IOI did not differ between males and females (B_ix_) ISI did not differ between males and females. All data are represented as mean ± SEM. *p < 0.05; **p < 0.01; ***p < 0.001

We examined the nose-poke distributions for both InSeq and OutSeq trials (Figures 3Av and 3Avi). The InSeq distribution showed the proportion of nose-pokes in males and females remained similar. On OutSeq trials, however, we see a modest difference between male and female poke times. Females tended to poke for OutSeq trials close to the 1s threshold (~0.8 –0.99s) whereas males continued to show short pokes (~0.1 – 0.4s) in both InSeq and OutSeq trials. We evaluated whether males and females differed in nose-poke times on InSeq_correct_ and OutSeq_correct_ trials (Figure 3Avii). No significant differences were detected between males and females for InSeq_correct_ trials (t_(373)_ = 1.195 *p* = 0.233). However, in OutSeq_correct_ trials we found a significant difference, with females showing a slightly longer hold time (t_(373)_ = 9.609, *p* = 1.119 x 10^-19^; Male: 0.446 ± 0.020; Female: 0.678 ± 0.014) which corresponds with what we observed in the poke distribution. We then looked at the time rats spent between each odor trial (inter-odor-interval; IOI) and ISI. The IOI did not differ significantly between males and females (Figure 3Aviii; t_(767)_ = 1.495, *p* = 0.135), suggesting rats collected water rewards and engaged with the odors at similar rates. Furthermore, we found no effect by group on the ISI (Figure 3Aix; t_(757)_ = 0.312, *p* = 0.755), suggesting rats in both groups alternated at similar rates between sequences.

Once rats reach criterion in the TwoOdor training stage, they are moved to ThreeOdors. In this stage, one more odor is added to each sequence (Figure 3Bi; Seq1: A_1_B_1_C_1_ and Seq2: A_2_B_2_C_2_). Rats were trained to poke three times on either side with at least ~65% accuracy before being introduced to the OutSeq probe trials. With this three-item sequence the number of possible OutSeq configurations has expanded, making this the first stage to test sequence memory. Rats were trained up until they demonstrated sufficient sequence memory (≥ 0.2 SMI) for three consecutive trials to move onto the final training stage. We averaged the first twenty sessions to measure learning. We found a significant difference in the first five sessions where males seemed to learn at a faster rate compared to females (Figure 3Bii; Sessions_1-5_: t_(16)_ = −2.471, *p* = 0.025). However, this difference diminished with the subsequent sessions (Sessions_5-10_: t_(14)_ = 1.131, *p* = 0.277; Sessions_10-15_: t_(12)_ = −0.037, *p* = 0.971; Sessions_15-20_: t_(9)_ = −0.206, *p* = 0.842).

We then examined the overall SMI using data from after subjects had reached the behavioral criterion. We found no significant differences between males and females (Figure 3Biii; t_(52)_ = −1.755, *p* = 0.085, Male: 0.428 ± 0.015, Female: 0.394 ± 0.012). We also observed no significant interactions between sex and sequence (F_(3,104)_ = 0.059, *p* = 0.808). A simple main effect analysis showed that sex and sequence did not have a statistically significant effect on SMI (Sex: *p* = 0.094; Sequence: *p* = 0.182).

We then evaluated the nose-poke distributions for both InSeq and OutSeq trials (Figure 3Bv and 3Bvi). The InSeq distribution shows a similar proportion of nose-pokes in males and females. The OutSeq trials showed that males poked between ~0.4 - 0.7s whereas females tended to poke between ~0.8 – 0.95s. We also saw more short pokes from males in both InSeq and OutSeq trials. We evaluated non-mnemonic behaviors by examining nose-poke behaviors, IOI, and ISI. Additionally, InSeq_correct_ nose-poke times between males and females did not significantly differ (Figure 3Bvii: t_(344)_ = 1.718, *p* = 0.087). There was a significant difference in OutSeq_correct_ trials with females holding longer (Figure 3Bvii; t_(344)_ = 5.619, *p* = 3.965 x 10^-8^; Males: 0.501 ± 0.021; Females: 0.643 ± 0.015). The IOI and ISI did not differ significantly between males and females (Figure 3Bviii and 3Bix; IOI: t_(562)_ = 1.568, *p* = 0.118; ISI: t_(562)_ = 1.431, *p* = 0.153). Overall, these results suggest that males and females do not differ in working memory. However, males show a slight advantage in learning the OutSeq rule during the ThreeOdor training stage.

Finally, upon successfully completing the ThreeOdor training stage, an additional odor is introduced to each sequence (Seq1: A_1_B_1_C_1_D1 and Seq2: A_2_B_2_C_2_D_2_). Rats were trained until they achieved ~65% accuracy in differentiating between InSeq and OutSeq (≥ 0.2 SMI). Once this criterion was met, we then ran the rats for 15 consecutive sessions (Figure 4). We took the averaged results from 25 sessions after the task was learned in order to describe overall learning and performance. We found no significant difference in learning between males and females. (Figure 4B; Sessions_1-5_: t_(16)_ = 0.221, *p* = 0.828; Sessions_5-10_: t_(16)_ = −0.176, *p* = 0.862; Sessions_10-15_: t_(16)_ = −0.214, *p* = 0.833; Sessions_15-20_: t_(16)_ = −0.437, *p* = 0.668; Sessions_20-25_: t_(14)_ = 1.701, *p* = 0.111). By session ten, most rats reached behavioral criterion (asymptotic sequence memory performance levels over multiple sessions). Overall, rats demonstrated strong sequence memory (Figure 4C; Male: 0.262 ± 8.690 x 10^-3^, Female: 0.272 ± 8.120 x 10^-3^) and performance did not differ significantly between males and females (Figure 4Ci; SMI: t_(268)_ = 0.784, *p* = 0.434). Both males and females performed well above chance levels on sequence 1 (Male: 0.268 ± 0.017; Female: 0.280 ± 0.018) and sequence 2 (Male: 0.279 ± 0.016; Female: 0.318 ± 0.018), with no significant interactions between sequences or sex (Figure 4Cii; F_(1,479)_ = 0.608, *p* = 0.436). Simple main effect analysis showed that sex and sequence did not have a statistically significant effect on SMI (Sex: *p* = 0.160; Sequence: *p* = 0.177), indicating rats successfully switched between the two sequences.

**Figure 4.**
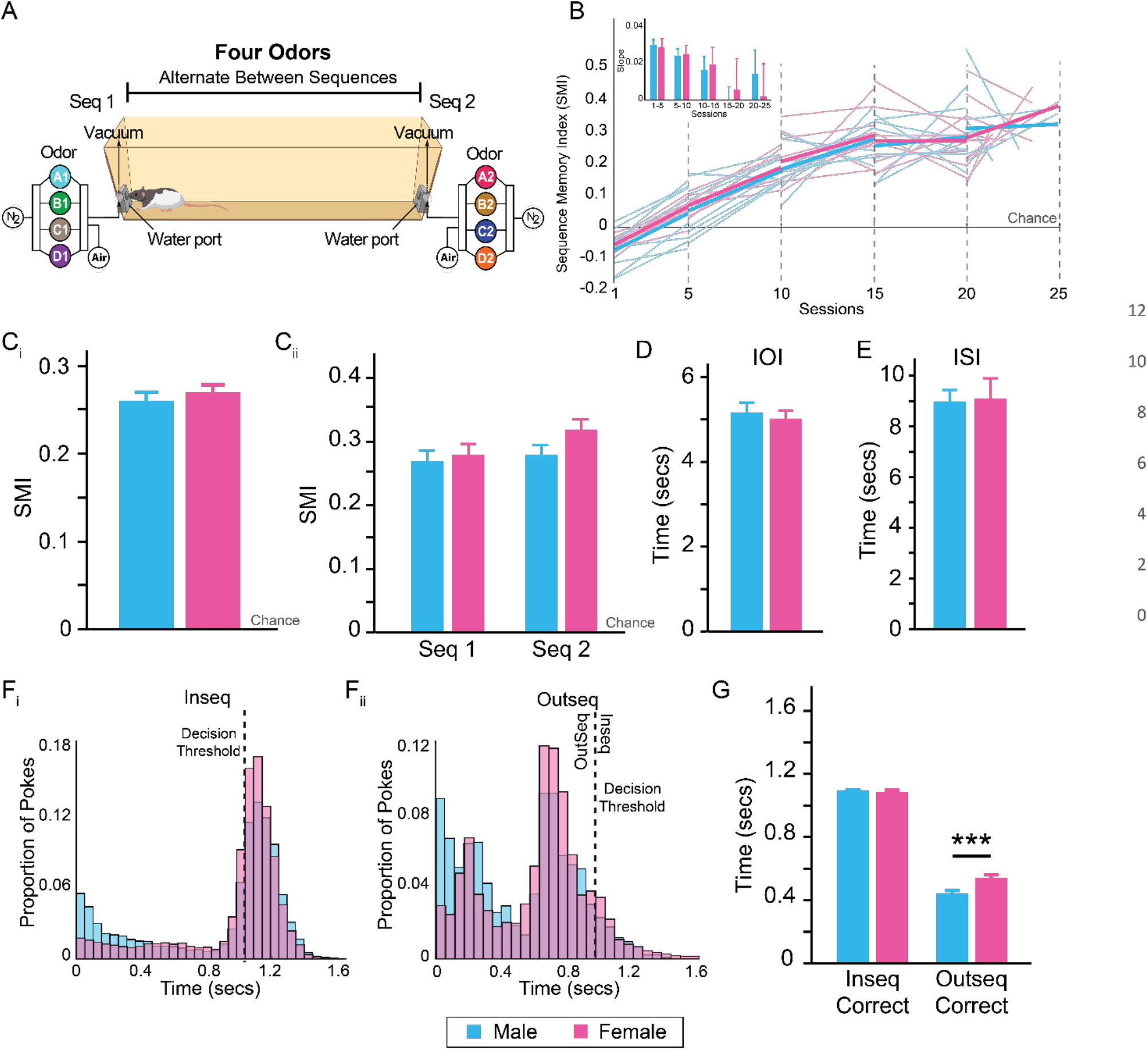
Males and Females Perform Similarly on the Sequence Memory Task. (A) Rats were trained train in the final stage of the sequence memory task with four odors in each sequence (Seq1: A_1_B_1_C_1_D_1_ and Seq2: A_2_B_2_C_2_D_2_). (B) Males and females showed no significant differences in the rate of learning across sessions. (C_i_) SMI was not significantly different between males and females. (C_ii_) SMI did not show any interaction effects between sequence 1 and sequence 2 and sex. (D) IOI was not statistically different between males and females. (E) ISI was not statistically different between males and females. (F_i_) Male and female poke distributions are relatively similar for InSeq trials (F_ii_) OutSeq poke distributions show subtle changes with males demonstrating an increase in shorter pokes compared to females. (G) InSeq_correct_ trials did not differ between males and females. OutSeq_correct_ trials showed a significant difference between sexes. All data are represented as mean ± SEM. *p < 0.05; **p < 0.01; ***p < 0.001

We evaluated non-mnemonic effect of sex by examining IOI and ISI. The IOI and ISI did not differ significantly between males and females (Figure 4D and 4E; IOI: t_(278)_ = −0.569, *p* = 0.570; ISI: t_(278)_ = 0.162, *p* = 0.871). We analyzed the poke distribution between males and females (Figure 4Fi and Fii). The InSeq and OutSeq distribution showed the proportion of nose-pokes remained similar between males and females. However, there was a slight increase in the proportion of InSeq and OutSeq nose-pokes near the short distribution peak for males (~0.1 – 0.2s). We also examined whether sex affected nose-poke times on InSeq_correct_ and OutSeq_correct_ trials (Figure 4G). No significant differences were detected in InSeq_correct_ trials (t_(268)_ = −1.925, *p* = 0.055). In the OutSeq_correct_ trials there was a significant increase in nose-poke time for females (t_(268)_ = 3.617, *p* = 3.560 x 10^-4^; Male: 0.443 ± 0.020; Female: 0.544 ± 0.019). Overall, these results demonstrate that males and females do not differ in their overall sequence memory. However, there are slight differences in their hold times for OutSeq trials, which may indicate uncertainty regarding whether a trial was InSeq or OutSeq rather than a deficit related to basic nose-poke behavior or sequence memory.

### Estrous Cycle in Sequence Memory

Generally, the results demonstrate that males and females do not differ in learning and performance on the sequence memory task. We then looked at estrous cycle phases within females, as the fluctuations of female hormone levels are important factors to consider when working with female animals. Figure 5A illustrates the hormone level in female rats during each of the four phases of the estrous cycle. In the proestrus phase, ovulation begins with the sharp increase and decrease of progesterone, increase of estradiol, and decrease of luteinizing hormone (LH). In the estrus phase, known as the “heat phase”, progesterone remains steady, estradiol peaks and begins to decrease while LH decreases. In the metestrus phase, all hormones remain steady. Finally, in the diestrus phase LH begins to increase slowly.

**Figure 5.**
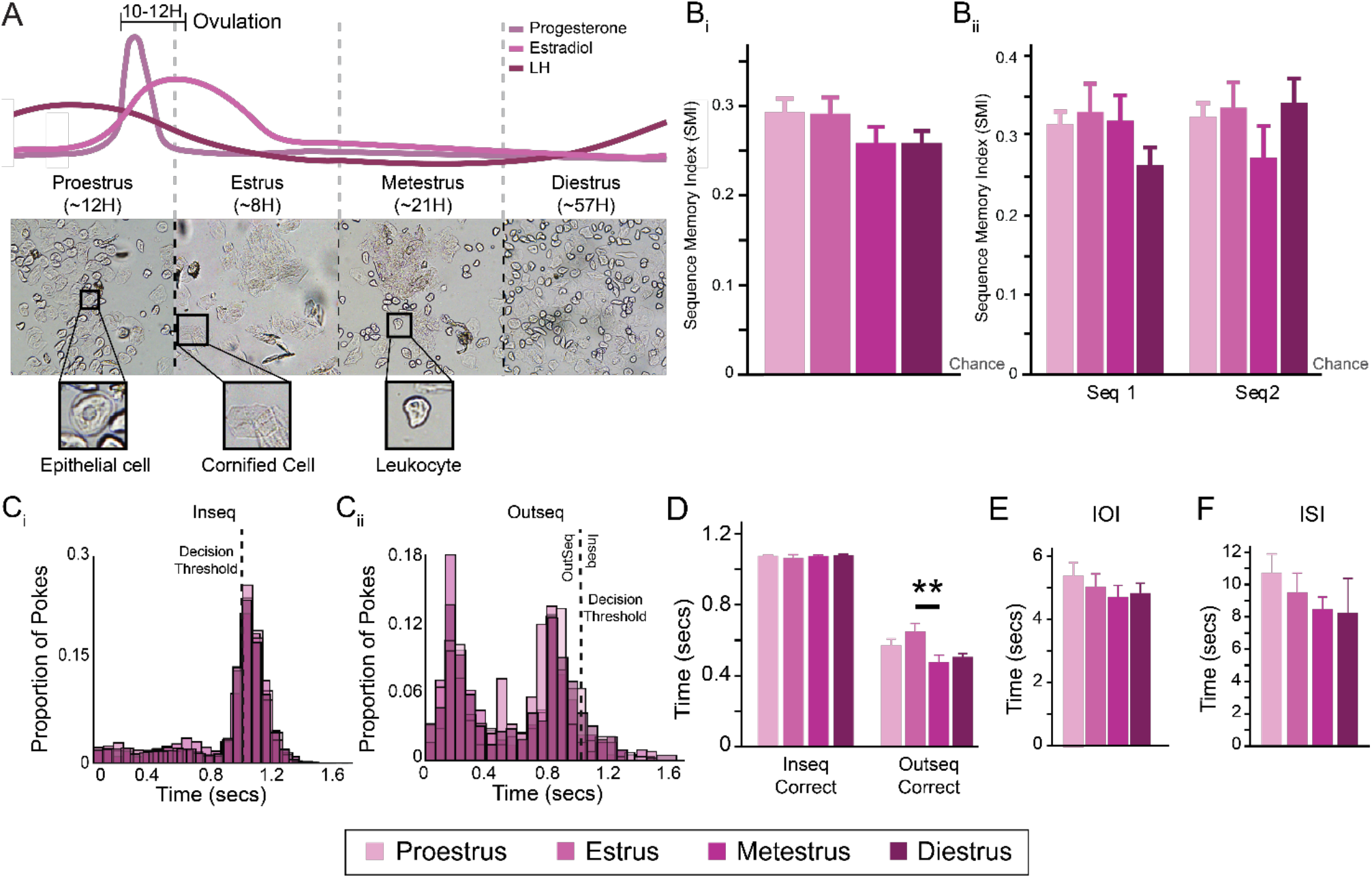
Estrous Cycle Does Not Influence Sequence Memory. (A) Rat ovulation hormone level fluctuations are shown for each stage as well as the amount of time each stage lasts. Lavage sample images of each phase of the estrous cycle with zoomed images of the primary cell types (adapted from Goldman et al., 2007). (B_i_) SMI was not different between estrous cycle phases. (B_ii_) SMI for sequence 1 and sequence 2 showed no significant differences for the estrous cycle phases (C_i_) InSeq poke distributions were relatively similar between estrous cycle phases. (C_ii_) OutSeq poke distributions showed subtle behavioral differences with estrus showing slightly increased short pokes. (D) InSeq_correct_ trials did not differ between estrous cycle phases. OutSeq_correct_ poke times were significantly different between estrus and metestrus. (E) IOI was not statistically different between estrous cycle phases. (F) ISI was not statistically different between estrous cycle phases. All data are represented as mean ± SEM. *p < 0.05; **p < 0.01; ***p < 0.001

We first examined the overall SMI once rats reach asymptotic levels on the fully learned sequence memory task. We found no significant differences between the estrous cycle phases (Figure 5Bi; F _(3, 146)_ = 1.405, *p* = 0.244). We also observed no significant differences between estrous and sequence (Figure 5Bii: F_(7, 270)_ = 3.56, *p* = 0.078). Simple main effect analysis showed that estrous and sequence were not statistically significant (Estrous: *p* = 0.433; Sequence: *p* = 0.322). We then observed the general poke distributions for InSeq and OutSeq trials between the estrous cycle phases. For InSeq trials, all estrous cycle phases showcased a cluster of pokes slightly after the 1 s decision threshold (Figure 5Ci). For OutSeq trials, there are two main peaks of nose-pokes around 0.2 s and 0.8 s (Figure 5Cii). We did observe that there was a slight increase in poke times during estrus (~0.2 – 0.3s) in comparison to the other three phases. Next, we evaluated the nose-poke times for InSeq and OutSeq correct trials. There were no significant differences in InSeq_correct_ trials between the estrous cycle phases (Figure 5D; F_(3, 149)_ = 0.667, *p* = 0.573). There was a significant difference in OutSeq_correct_ trials (Figure 5D; F_(3, 149)_ = 3.301, *p* = 0.022). Post hoc comparisons using Bonferroni test revealed a significant difference between estrus and metestrus (*p* = 0.033) with a mean difference poke time of 0.177s. We explored non-mnemonic effects relevant to the sequence task. We looked at IOI and ISI and found no significant differences between the estrous cycle phases within those conditions (Figure 5E and 5F: IOI: F_(3, 149)_ = 0.676, *p* =0.568; ISI: F_(3, 149)_ = 0.416, *p* = 0.742). These results that the estrous cycle does not strongly affect overall sequence memory performance.

## DISCUSSION

### Summary of Main Findings

We evaluated the role of sex in rats during training and testing phases of a sequence memory task. Although this task has been used in previous studies (Allen et al., 2014, 2016a; Jayachandran et al., 2019). here we describe the stages of training and its performance criterion for the first time. Overall, both male and female rats learned the sequence memory task at relatively the same rate. However, there were a few differences between males and females. Males tended to have a modest number of shorter poke times compared to females, which was more evident once OutSeq trials were introduced, but this did not affect overall accuracy. Moreover, males seemed to learn at a somewhat faster rate once the sequence was expanded to include more than one OutSeq position (ThreeOdor) but this effect did not extend to the full sequences. In general, there were no noticeable sex differences in sequence memory once the task was fully learned. This finding alone, however, did not address whether the lack of sex differences could have been confounded by female rats’ estrous cycle. Therefore, we looked at estrous cycle specifically and found that regardless of phase, females performed similarly well on the sequence memory task.

### No Evidence that Sequence Memory Differs in Males and Females

To investigate sex differences in sequence memory, we examined males and females at each training stage of the sequence task. One main difference we observed was with the introduction of the three-item sequence. At this stage, we found males had a faster learning curve compared to females. This advantage could be due to males’ tendency for short pokes, allowing them more opportunities to have chance correct responses for OutSeq trials and thus providing a modest boost in discovering the rules. This may not reflect sex differences in learning per se, but rather sex differences in strategies (Gruene et al., 2015). This has been suggested before for spatial learning paradigms, in which male rats outperform females by using a more direct strategy (McCarthy & Konkle, 2005). This difference could also be due to Type I family-wise error. As multiple tests were run at multiple stages and therefore these differences could be minor. Moreover, this advantage was not observed once the rats moved to the final stage of training (FourOdors), where overall sequence memory did not differ between sexes. This is further supported by Reeders et al., (2021) who did not observe sex differences in humans performing an analogous task. Importantly, sequence memory effects were consistent across two different sequences. This eliminated the possibility that males and females show preference for a specific sequence or odors throughout the entirety of a session.

Apart from examining the general learning rates at each stage, we also performed several analyses to test whether males and females differed in non-memory related behaviors in the sequence task. We saw no effects between males and females on reward retrieval activity (ITI & IOI), on the time it took to run between sequence (ISI), or on the overall frequency of nose pokes in which rats held the nose poke response for ≥ 1 s (InSeq trials). We did observe females had modestly longer hold times compared to males for OutSeq trials. This was more clearly revealed in the analysis of poke time histogram where males showed a tendency to have shorter poke times whereas females showed a tendency to have longer poke times.

Although, several studies have shown a robust male advantage in working and reference memory (Roof et al., 1993; Veng et al., 2003), other studies indicate no differences or a female advantage (Bucci et al., 2021; Healy et al., 1999; Lamberty & Gower, 1988). These confounding observations in sex differences have been associated with different factors within the task parameters. For example, a consistent start position for each trial has been associated with a lack of sex differences (Roof & Stein, 1999). Pre-training has also been shown to reduce sex differences (Jonasson, 2005a; Perrot-Sinal et al., 1996). Generally speaking, male and female rodents typically reach identical levels of performance on most cognitive tasks although their rates and strategies of learning may differ.

### No Evidence Sequence Memory Differs Across Estrous Cycle

Several studies have postulated that hormone fluctuations across the estrus cycle may account for sex differences (McEwen & Milner, 2017; Sherry & Hampson, 1997; Warren & Juraska, 1997). However, no clear consensus has emerged regarding the influence of estrous cycle on learning and memory. It has been argued that estrous cycle cannot be causally linked to more variability in females relative to males (Prendergast et al., 2014). Pompili et al. (2010) observed no variation in performance associated with cycle phase when using the radial-arm maze. Stackman et al. (1997), using a delayed nonmatching-to-sample paradigm in the radial-arm maze, found no effect of estrous cycle. Nevertheless, it is important to consider sex hormones in behavior. Therefore, we examined sequence memory across phases of estrous cycle. In general, we did not find differences between the phases of estrous cycle in sequence memory. It is possible that pre-training might have reduced the sensitivity of this assessment (Bannerman et al., 1995; D. Saucier & Cain, 1995). However, effects of the cycle on memory are somewhat subtle and inconsistent, so they may wash out when averaging across multiple females.

Interestingly, while sequence memory was not affected by the day of estrous cycle, performance on estrus was characterized by longer OutSeq poke times. One natural issue presented by measuring estrous cycle for this task is the asymmetry of the length of stages; because each stage averages a different length of time, we would expect to see far more trials occurring during these longer stages than the shorter ones. To the extent that proestrus, metestrus and diestrus is over-represented compared to estrus, which could have skewed the group mean. Several studies report that spatial memory tested in the Morris water maze or object placement tasks was enhanced during proestrus relative to estrus and/or diestrus in mice and rats (Frick & Berger-Sweeney, 2001; Frye, 1995; Paris & Frye, 2008; Pompili et al., 2010). However, the reported proestrus advantage in spatial tasks is inconsistent with other data in rats showing enhanced spatial reference memory during estrus relative to proestrus (Frye, 1995; Sutcliffe et al., 2007a; Warren & Juraska, 1997) or no detectable effect of the cycle on memory (Berry et al., 1997; Bucci et al., 2008; Cimadevilla et al., 2000; Conrad et al., 2004; Pompili et al., 2010; Stackman et al., 1997). While the exact cognitive bases are not yet known, Korol & Kolo (2002) have suggested that the direction of estrogen effects on performance depends upon the availability of strategies or solutions that match the participation of neural or cognitive systems. Thus, the general effects of estrogen on memory are somewhat ambiguous (Dohanich, 2002). The sequence task used here is known to involve multiple strategies including temporal contexts, ordinal representations and working memory (e.g., Reeders et al, 2021; Jayachandran et al., 2019)

### Conclusions

We present evidence that sex and estrous cycle do not influence memory for sequences of events suggesting that this core aspect of episodic memory does not differ between the sexes. These data are consistent with few other studies that have investigated sex differences and estrous cycle as it pertains to behavior in rodents as well as in humans (Bucci et al., 2021; Healy et al., 1999; Reeders et al., 2021; Schmidt et al., 2009; Stackman et al., 1997). However, further studies are needed to fully characterize the effects of sex hormones on sequence memory in both male and female rats by comparing the effects of gonadectomy and possible hormone replacement. For now, the consideration of sex and estrous cycle as a biological variable will have significantly advanced our understanding of the basic mechanisms of learning and memory.

## ACKNOWLEDGMENTS

This work was supported by NIH grants R01 MH113626 and F99 NS119001. A special thanks to all the members of the Allen Lab, specifically our undergraduate research assistants, Andy Garcia, Sofia Levya, and Chiara Pavon, who helped with task training and data collection. Also, we would like to thank the Animal Care Facility and Dr. H. Vinerean DVM, DACLAM.

## AUTHOR CONTRIBUTIONS

Conceptualization, T.A.A. and M.J.; Methodology, T.A.A. and M.J.; Investigation, M.J., P.L., F.P.R.; Writing – Original Draft, M.J., P.L., T.A.A.; Writing – Review and Editing, M.J., P.L., F.P.R., T.A.A., R.P.V.; Funding Acquisition, M.J., T.A.A. and R.P.V.; Resources, T.A.A., R.P.V.; Supervision, T.A.A.

